# Unravelling the impact of TLR4 and sex on chronic alcohol consumption-induced lipidome dysregulation in extracellular vesicles

**DOI:** 10.1101/2024.09.05.611414

**Authors:** Carla Perpiñá-Clérigues, Susana Mellado, Cristina Galiana-Roselló, Francisco García-García, María Pascual

## Abstract

The lipids that form extracellular vesicles (EVs) play critical structural and regulatory roles, and cutting-edge bioinformatics strategies have shown the ability to decipher lipid metabolism and related molecular mechanisms. We previously demonstrated that alcohol abuse induces an inflammatory immune response through Toll-like receptor 4 (TLR4), leading to structural and cognitive dysfunction. This study evaluated how TLR4 and sex as a variable (male/female) impact the lipidome of plasma-resident EVs after chronic alcohol exposure. Using a mouse model of chronic ethanol exposure in wild-type and TLR4-deficient mice, enrichment networks generated by LINEX^2^ highlighted significant ethanol-induced changes in the EV lipid substrate-product of enzyme reactions associated with glycerophospholipid metabolism. We also demonstrated ethanol-induced differences in Lipid Ontology enrichment analysis in EVs, focusing on terms related to lipid bilayer properties. A lipid abundance analysis revealed higher amounts of significant lipid subclasses in all experimental comparisons associated with inflammatory responses and EV biogenesis/secretion. These findings suggest that interrogating EV lipid abundance with a sensitive lipidomic-based strategy can provide deep insight into the molecular mechanisms underlying biological processes associated with sex, alcohol consumption, and TLR4 immune responses and open new avenues for biomarker identification and therapeutic development.

## Introduction

Lipids represent crucial components of cell membranes and participate in a range of cellular functions. Current studies indicate that the lipid content of extracellular vesicles (EVs) plays critical structural and regulatory roles during EV biogenesis, release, targeting, and cellular uptake ^1^. These nanovesicles act as important intercellular communicators and it has been recently demonstrated that their lipid content is involved as diagnostic biomarkers in pathological processes ^2,3^. Mass spectrometry-based lipidomics combined with dedicated computational tools represents a powerful means of identifying and quantifying lipids in cells, tissues, or bodily fluids ^4^. Recently, the lipid network explorer (LINEX^2^) emerged as a novel bioinformatics tool to decipher lipid metabolism and related molecular mechanisms by combining lipid classes and metabolic reactions ^5^. These methodological developments allowed the identification of novel lipid targets and the discovery of novel clinical biomarkers ^6,7^.

Alcohol is a neurotoxic compound that alters brain structure and function by inducing alcohol dependence and prompting the development of behavioral, cognitive, and psychiatric disorders ^8^. Although the exact molecular mechanisms at play remain incompletely understood, our findings have revealed the participation of innate immune response through the activation of the Toll-like receptor 4 (TLR4) immune signaling pathway in glial cells ^9,10^ and the brain in vivo ^11,12^. Activated TLR4 signaling during chronic alcohol exposure triggers the induction of brain inflammatory cytokines, gliosis, and the loss of myelin ^13^ and causes brain damage and long-term cognitive dysfunction ^14^. TLR4 knockout in mouse models (TLR4-KO) protects against ethanol-induced inflammatory response, brain damage, and cognitive dysfunction ^15,16^. In addition to the immune response, sex also represents a biological variable to consider in alcohol addiction and treatment ^17,18^; moreover, sex plays a significant role in the observed differences between immune responses and lipid metabolism in male and female humans ^19^ and rodents ^20^.

Considering that our recent studies reported the enrichment of specific lipid species in plasma EVs of human females associated with inflammatory immune responses ^21,22^, we have now employed a sensitive lipidomic-based strategy to demonstrated the involvement of TLR4 and sex differences in the alterations of EV lipid metabolic networks and lipid abundance induced by chronic ethanol exposure in female and male mice.

## Experimental Section

### Animals and alcohol exposure

Female (F) and male (M) wild type (WT, TLR4^+/+^) (Harlan Ibérica S.L., Barcelona, Spain) and TLR4 knockout (TLR4-KO, TLR4^-/-^, KO) mice, kindly provided by Dr S. Akira (Osaka University, Osaka, Japan) with C57BL/6 genetic backgrounds, were used in this study. Animals were kept under controlled light and dark conditions (12 h/12 h) at 23°C and 60% humidity. Animal experiments were conducted in accordance with the guidelines set out in the European Community Council Directive (2010/63/ECC) and Spanish Royal Decree 53/2013, modified by Spanish Royal Decree 1386/2018, and were approved by the Ethical Committee of Animal Experimentation of the CIPF (Valencia, Spain).

Two-month-old animals were housed (2–4 animals/cage) and divided into eight groups (n=6 mice/group): female and male WT mice for control and chronic ethanol exposure and female and male TLR4-KO mice for control and chronic ethanol exposure. Mice were provided with water (control) or water containing 10% (v/v) ethanol for three months, during which time they were fed a solid diet *ad libitum*. Daily food and fluid intake in the four groups were carefully measured; no differences were previously demonstrated ^12,23^. Similarly, blood ethanol levels remained comparable in the ethanol-exposed groups (WT and TLR4-KO) (125 ± 20 mg/dL) ^12,23^. Animals were anesthetized at the end of ethanol exposure, and whole blood samples were collected from the hepatic portal vein. After centrifugation, the separated plasma was snap-frozen in liquid nitrogen and stored at -80°C until use. **Figure 1A** summarizes the experimental groups.

**Figure 1.**
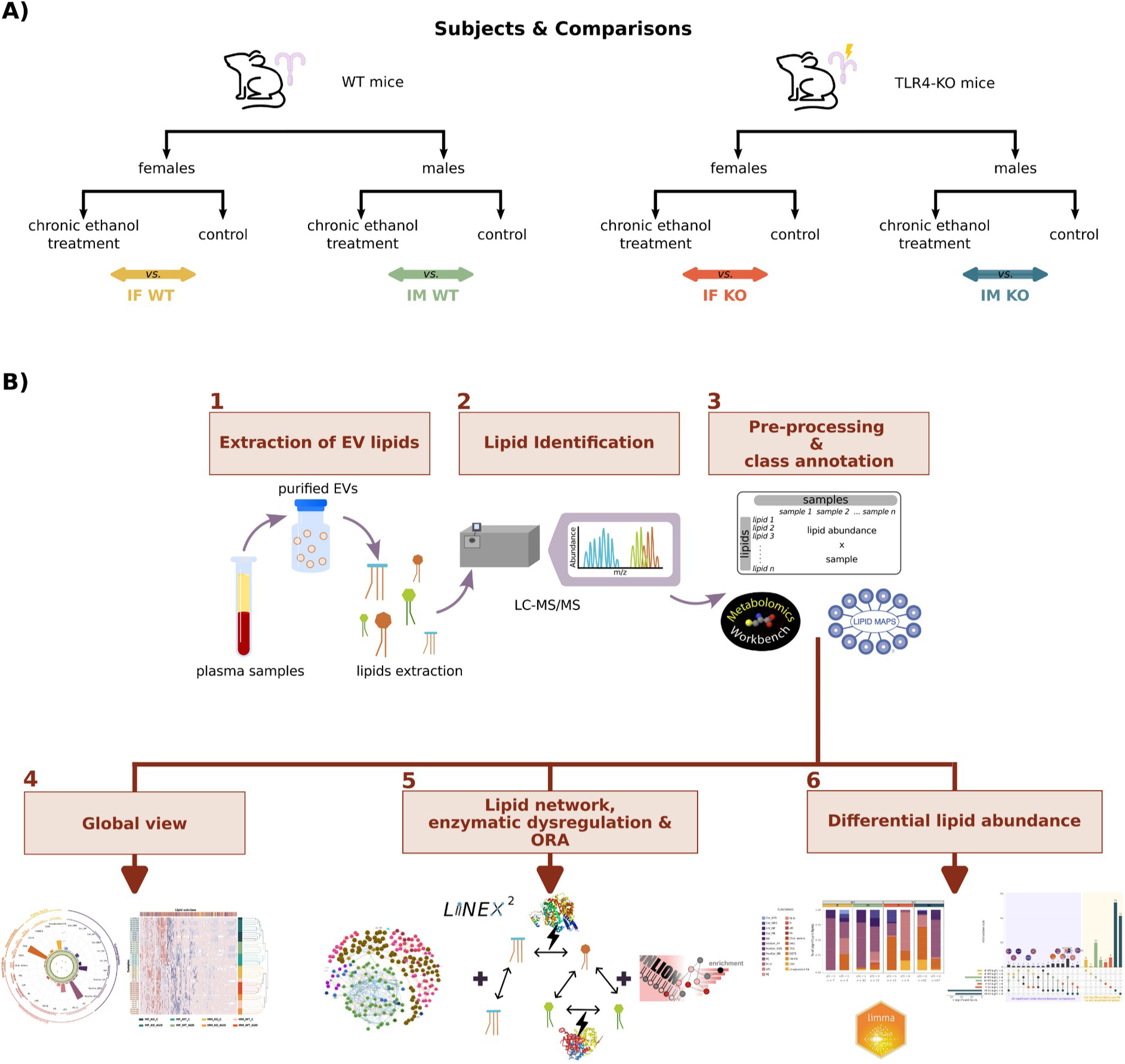
Experimental design and lipidomic workflow. **(A)** Description of the experimental groups and the comparisons made. Ethanol exposure vs. control in female WT mice (IF WT), ethanol exposure vs. control in male WT mice (IM WT), ethanol exposure vs. control in female TLR4-KO mice (IF KO), and ethanol exposure vs. control in male TLR4-KO mice (IM KO). **(B)** Lipids were extracted for quantification and identification using liquid chromatography with tandem mass spectrometry (LC-MS/MS) after isolating extracellular vesicles (EVs) from mouse plasma in the lipidomic workflow. Following data normalization, lipid class annotation, and exploratory analyses, the LINEX^2^ platform provided a global reaction network, the subgraph with the most significant average substrate-product change, and a list of target lipids for further enrichment analysis using LION-web. Lastly, differential analyses assessed lipid abundance. Abbreviations: ORA – over-representation analysis.

### Isolation of extracellular vesicles from mouse plasma

Plasma EVs were isolated by a total exosome isolation kit (catalog number 4484450, Invitrogen, USA), following the manufacturer’s instructions. 250 μL of mouse plasma was used to isolate EVs, which were collected and frozen at -80°C until processing.

### Extracellular vesicle characterization by transmission electron microscopy and nanoparticle tracking analysis

Freshly isolated EVs were fixed with 2% paraformaldehyde and prepared as previously described (Ibáñez et al., 2019). Preparations were examined using an FEI Tecnai G2 Spirit transmission electron microscope (TEM; FEI Europe, Eindhoven, the Netherlands) with a Morada digital camera (Olympus Soft Image Solutions GmbH, Münster, Germany). In addition, the absolute size range and concentration of EVs were analyzed using NanoSight NS300 Malvern (NanoSight Ltd., Minton Park, UK) as previously described ^24^.

### Lipid extraction

Lipids were extracted from equal amounts of plasma EVs (0.2 ml/sample) using a modified Folch extraction procedure. The last phase containing the lipids was transferred to fresh tubes, dry vacuumed with nitrogen, and stored at -80°C until further analysis. Dried samples were resuspended with isopropanol for different liquid chromatography-mass spectrometry (LC-MS) acquisition methods (positive and negative ion modes).

### Liquid chromatography with tandem mass spectrometry analysis

In fully automated quadrupole time-of-flight (QToF) acquisition mode, a pooled human lipid extract representing the 36 samples (4 conditions × 9 replicates) was acquired by iterative tandem mass spectrometry (MS/MS). Experimental methods for chromatography and auto MS/MS mass spectrometry were used as previously described ^25,26^ with minor modifications. Briefly, sample separation was performed using an Agilent 1290 Infinity LC system coupled to the 6550 Accurate-Mass QTOF (Agilent Technologies, Santa Clara, CA, USA) with electrospray interface (Jet Stream Technology) operating in positive-ion (3500 V) or negative-ion (3000 V) mode and high sensitivity mode. The optimal conditions for the electrospray interface were a gas temperature of 200 °C, drying gas of 12 L/min, nebulizer of 50 psi, sheath gas temperature of 300 °C, and sheath gas flow of 12 L/min. Lipids were separated on an Infinity Lab Poroshell 120 EC-C18 column (3.0 ×100 mm, 2.7 μm) (Agilent, Santa Clara, CA, USA). Under optimized conditions, the mobile phase consisted of solvent A (10 mM ammonium acetate and 0.2 mM ammonium fluoride in 9:1 water/methanol) and solvent B (10 mM ammonium acetate and 0.2 mM ammonium fluoride in 2:3:5 acetonitrile/methanol/isopropanol) using the following gradient: 0 min 70%B, 1 min 70%B, 3.50 min 86%B, 10 min 86%B, 11 min 100%B, 17 min 100%B operating at 50 °C and a constant flow rate of 0.6 mL/min. The injection volume was 5 µL for positive and negative modes.

The Agilent MassHunter Workstation Software was employed for data acquisition. LC/MS Data Acquisition B.10.1 (Build 10.1.48) was operated in auto MS/MS mode, and the three most intense ions (charge states, 1-2) within 300–1700 m/z mass range (over a threshold of 5000 counts and 0.001%) were selected for MS/MS analysis. The quadrupole was set to a "narrow" resolution (1.3 m/z), and MS/MS spectra (50–1700 m/z) were acquired until 25,000 total counts or an accumulation time limit of 333 ms. To assure the desired mass accuracy of recorded ions, continuous internal calibration was performed during analyses using signals m/z 121.050873 and m/z 922.009798 for positive mode and signals m/z 119.03632 and m/z 980.016375 for negative mode. Additionally, all-ions MS/MS data ^27^ were acquired on individual samples, with an MS acquisition rate of three spectra/second and four scan segments (0, 10, 20, and 40 eV).

### Lipid annotator database

Five sets of five iterative MS/MS data files from pooled mice lipid extracts were analyzed with Lipid Annotator software 1 as the first step in the lipidomics workflow. This study used a novel software tool (Lipid Annotator) ^28^ with a combination of Bayesian scoring, a probability density algorithm, and non-negative least-squares fit to search a theoretical lipid library (modified LipidBlast) developed by Kind et al. ^29,30^ to annotate the MS/MS spectra.

Agilent MassHunter Lipid Annotator Version 1.0 was used for all other data analyses. Default method parameters were used; however, only [M+H]^+^ and [M+NH4]^+^ precursors were considered for positive ion mode analysis, and only [M-H]^–^ and [M+HAc-H]^–^ precursors were considered for negative ion mode analysis. Agilent MassHunter Personal Compound Database and Library (PCDL) Manager Version B.08 SP1 was used to manage and edit the exported annotations.

### Lipid identification

The lipid Personal Compound Database and Library (PCDL) databases created were used for Batch Targeted Feature Extraction in Agilent MassHunter Qualitative version 10.0 on the respective batches of 36 all-ions MS/MS data files. The provided "Profinder - Lipids.m" method was adapted in MassHunter Qualitative software with modifications previously described by Sartain et al. ^26^. Data were analyzed using the Find by Formula (FbF) algorithm in MassHunter Qualitative Analysis. This approach uses a modified version of the FbF algorithm, which supports the all-ions MS/MS technique. Mass peaks in the low energy channel are first searched against the PCDL created for compounds with the same m/z values, and then a set of putative identifications is automatically compiled. For this list, the fragment ions in the MS/MS spectra from the PCDL are compared to the ions detected in the high-energy channel to confirm the presence of the correct fragments. The precursors and products are extracted as ion chromatograms and evaluated using a coelution score. The software calculates a number for abundance, peak shape (symmetry), peak width, and retention time. The resulting compounds were reviewed in the MassHunter Qualitative version, and the features that were not qualified were manually removed. MassHunter Qualitative results and qualified features were exported as a .cef file.

### Bioinformatic analyses

The strategy applied for this study was based on a transcriptomic analysis workflow, including the specifics of the lipidomic data. All bioinformatics and statistical analysis were performed using R software v.4.1.2 ^31^. **Figure 1B** illustrates the lipidomics workflow strategy.

### Data preprocessing

Data preprocessing included filter entities, normalization of abundance lipid matrix, and exploratory analyses. MassHunter Qualitative results (.cef file) were imported into Mass Profiler Professional (MPP) (Agilent Technologies) for statistical analysis. Entities were filtered based on frequency, selecting those consistently present in all replicates of at least one treatment. A percentile shift normalization algorithm (75%) was used, and datasets were baselined to the median of all samples, including negative and positive ion modes. Normalized data were labeled according to negative and positive ion modes, and all data were consolidated into a single data frame. This step was followed by exploratory analysis using hierarchical clustering, principal component analysis (PCA), and box and whisker plots by samples and lipids to detect abundance patterns between samples and lipids and batch effects anomalous behavior in the data. At this point, anomalously-behaving samples and outliers (values that lie over 1.5 × interquartile range (IQR) below the first quartile (Q1) or above the third quartile (Q3) in the dataset) were excluded for presenting robust batch effects with a critical impact on differential abundance analysis.

### Class annotation

Class annotation was conducted using the *RefMet* database ^32^ and compared with the *LIPID MAPS* database ^33^. The classification is hierarchical. As an initial step in this division, lipids were divided into several principal categories ("super classes") containing distinct main classes and sub-classes of molecules, devising a standard manner of representing the chemical structures of individual lipids and their derivatives. **Table S1** describes the abbreviations employed.

### Lipid network

The Lipid Network Explorer platform (LINEX^2^, https://exbio.wzw.tum.de/linex/) was used for lipid metabolic network analysis to gain insights into the role of TLR4 and sex differences in the EV lipid metabolism of ethanol-exposed animals. For this purpose, single lipid species were considered as the sum or molecular species regardless of their retention time and ion mode acquisition. Therefore, before conducting the analysis, the lipid nomenclature was evaluated to ensure that most lipids in the study were included. This review was conducted using the MetaboAnalyst 5.0 platform ^34^ and the LipidLynxX Converter tool (http://www.lipidmaps.org/lipidlynxx/converter) ^35^. Additionally, a manual lipid-by-lipid revision was performed to ensure accuracy.

LINEX^2^ analysis provides several results. The global lipid species network provides qualitative associations between species based on defined reaction types. The subgraph with the most significant average substrate-product changes was obtained through a lipid network enrichment algorithm, which considered enzymatic multispecificity and generated hypotheses regarding enzymatic dysregulation. This algorithm consists of a local search approach that greedily examines a search space by iteratively testing local candidate solutions for the one with an optimal objective function. Candidate solutions are generated by applying one of three operations: node insertion, deletion, and substitution to the solution from the last iteration or a randomly selected subgraph in the first iteration. Lastly, LINEX^2^ provided a target lipids list derived from the lipid subgraph, which was utilized for an enrichment analysis using LION-web (http://www.lipidontology.com/) (LION - Lipid Ontology) ^36^. This enabled a more in-depth examination of the functional significance and potential biological implications of the identified lipid alterations.

### Differential lipid abundance

The limma R package compared lipid abundance levels between groups ^37^. p-values were adjusted using the Benjamini & Hochberg (BH) procedure ^38^, and significant lipids were considered when the BH-adjusted p-value ≤ 0.05.

### Comparisons

A comparison between ethanol-exposed and untreated mice was performed in both sexes (F: female and M: male) for the two genotypes of mice (**Fig. 1A**) in all groups (**Table 1**), resulting in the following comparisons, where IF and IM indicate the impact of the ethanol exposure in females and males, respectively:

- Ethanol exposure vs. control in female WT mice (IF WT)
- Ethanol exposure vs. control in male WT mice (IM WT)
- Ethanol exposure vs. control in female TLR4-KO mice (IF KO)
- Ethanol exposure vs. control in male TLR4-KO mice (IM KO)

**Table 1.** Different groups used in the study , categorized by sex, genotype and ethanol exposure.

| Group | Abbreviation |
| --- | --- |
| Female WT mice exposed to ethanol | F_WT_ET |
| Control female WT mice | F_WT_C |
| Male WT mice exposed to ethanol | M_WT_ET |
| Control male WT mice | M_WT_C |
| Female TLR4-KO mice exposed to ethanol | F_KO_ET |
| Control female TLR4-KO mice | F_KO_C |
| Male TLR4-KO mice exposed to ethanol | M_KO_ET |
| Control male TLR4-KO mice | M_KO_C |

The statistics used to measure the differential patterns were the logarithm of fold change (LFC) to quantify the effect of differential lipid abundance analysis. A positive statistical sign indicates a higher mean for the variable in the first element of the comparison, whereas a negative statistical sign indicates a higher mean value for the second element. In the study, the first element corresponds to ethanol exposure, and the second element is used to control.

## Results and discussion

### Data exploration insights into the impact of TLR4 and sex on chronic ethanol exposure in mice

First, we evaluate the role of TLR4 and sex in the altered plasma EV lipidome after chronic ethanol exposure in mice, we first characterized EVs using electron microscopy and nanoparticle tracking analysis (**Fig. 2**). The TEM study revealed that the nano-sized particles isolated from plasma displayed typical exosomal characteristics in terms of size and shape (∼100 nm in diameter; **Fig. 2A**). In addition, we analyzed the size distribution and concentration of secreted nano-sized particles using NanoSight; these results indicated that the highest peak of the secreted nano-sized particles fell within the 100-200 nm range, encompassing the size range for EVs (**Fig. 2B**).

**Figure 2.**
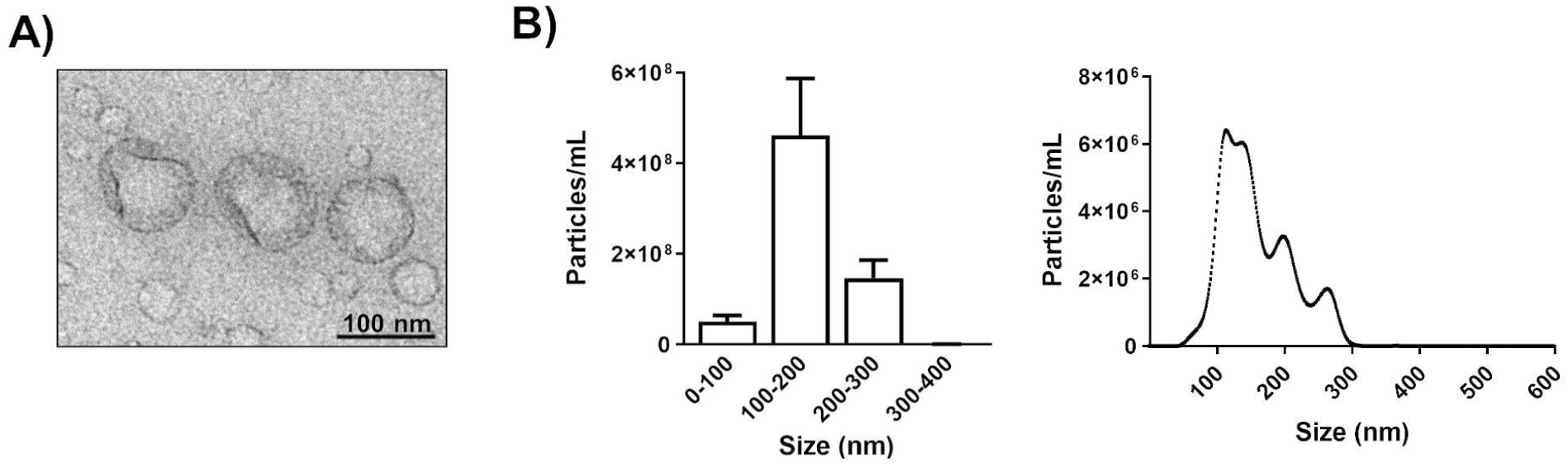
Characterization of extracellular vesicles isolated from mouse blood plasma. **(A)** Transmission electron microscopy image of mouse plasma extracellular vesicles (EVs). **(B)** Measurement of mouse plasma EV size distribution and concentration by nanoparticle tracking analysis.

A lipidomic analysis of all plasma EVs revealed 187 and 194 common lipid species across all groups using negative and positive ion modes, respectively. After normalizing sample data, we used the RefMet and LIPID MAPS databases to classify all lipids (381 species) into different subclasses and their upper levels (super and main classes) (**Fig. 3A** and **Table S2**). The sum of total lipid abundance by lipid subclass across the eight experimental groups revealed an enrichment of TAG and PC, and a moderate enrichment of SM, LPC, and Unsaturated FA subclasses in plasma EVs (**Fig. 3A**). The hierarchical clustering of EV lipid species, regardless of their subclass, revealed distinct lipid profiles for the eight experimental groups (**Table 1**); we identified lipidomic profiles based on sex, with samples of each sex grouping together (e.g., green and yellow-orange hues correspond to females and males, respectively) (**Figure 3B**). Additionally, we found that samples grouped by treatment and genotype (WT or TLR4-KO), especially in males and females, respectively (**Figure 3B**).

**Figure 3.**
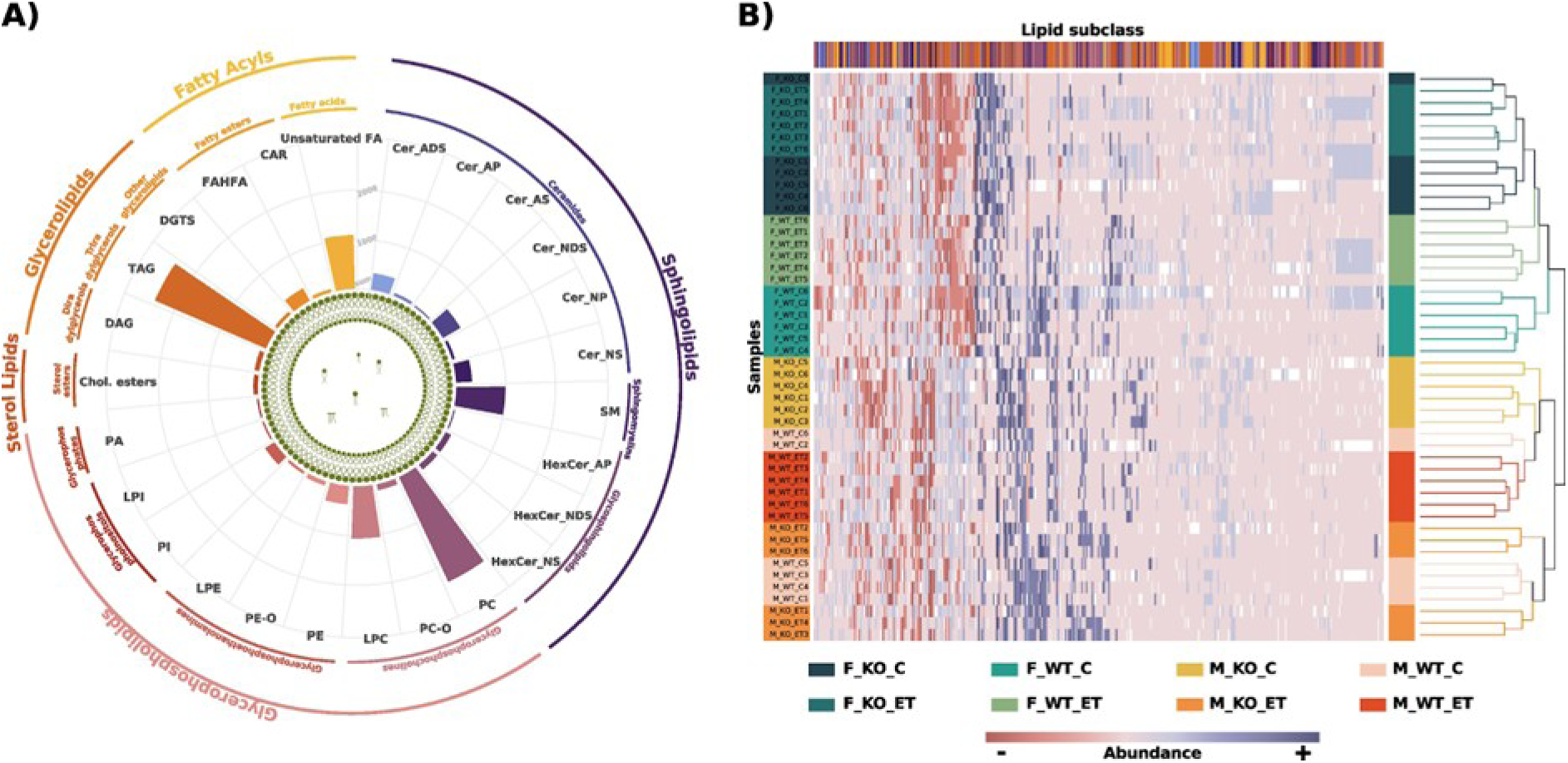
Lipid composition and distribution in extracellular vesicles isolated from the eight experimental groups. **(A)** The sum of total lipid abundance by lipid subclass across the eight experimental groups. Lipids were quantified as the log2 transformed value of the identified peak area by liquid chromatography with tandem mass spectrometry (LC-MS/MS). According to the RefMet classification, the inner and outer lines of the radar plots indicate the lipid main class and superclass, respectively. **(B)** Heatmap demonstrating the abundance patterns between subclass lipids (columns) and samples (rows). Lipid subclasses indicated by the same colors previously assigned in the radar plots. F: females, M: males, WT: wild-type, KO: TLR4-KO, C: control, ET: chronic ethanol exposure.

Overall, the data suggest that sex and genotype affect EV lipid distribution in control and alcohol-exposed mice, which agrees with several previous studies demonstrating the involvement of TLR4 activation and sex in the neuroinflammatory response induced by alcohol consumption ^11,13^.

### Lipid networks reveal glycerophospholipid metabolism alterations induced by chronic ethanol exposure

We aimed to determine the effect of TLR4 on lipid metabolism in chronically ethanol-exposed mice by analyzing the EV lipid content in males and females. For this purpose, we identified the most dysregulated lipid enzymes in the state comparisons using the LINEX ^2^ substrate-product ratio-based enrichment algorithm. **Figure 4A** reveals the global network of lipid species, which provides qualitative associations between species based on defined reaction types. We then obtained enriched subnetworks (reactions) with the most significant alteration in substrate-product between control and chronically ethanol-exposed male and female mice of both genotypes (WT and TLR4-KO) (**Figure 4B**). Interestingly, the resulting subnetworks comprised only PC and LPC lipid species in TLR4-KO male mice and WT females and males; meanwhile, TLR4-KO females displayed the additional LPE subclass (**Table S3**).

**Figure 4.**
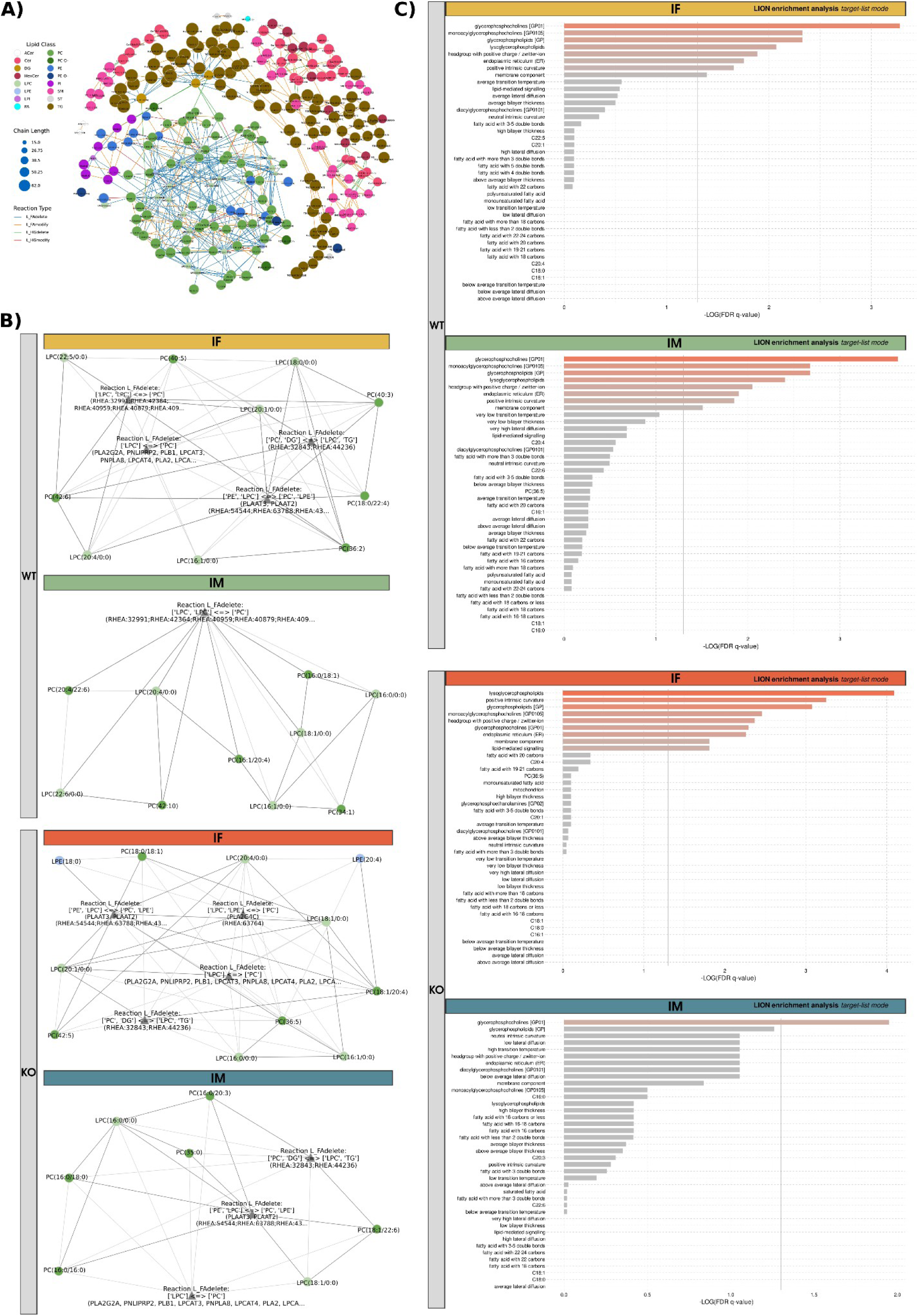
Lipid network analysis using LINEX^2^ in extracellular vesicles for the four study comparisons. **(A)** Global lipid network based on liquid chromatography with tandem mass spectrometry (LC-MS/MS) data from common lipids across different experimental groups. Colored spherical nodes depict lipid classes. Edge colors indicate the type of reaction connecting nodes. **(B)** Optimal subnetworks predicted by the enrichment algorithm for the following comparisons: ethanol exposure vs. control in female WT mice (IF WT), ethanol exposure vs. control in male WT mice (IM WT), ethanol exposure vs. control in female TLR4-KO mice (IF KO), and ethanol exposure vs. control in male TLR4-KO mice (IM KO). **(C)** Most enriched ontology terms result from using lipids in subnetworks as targets in the target list mode for the four comparisons in the ontology enrichment analysis in LION/web.

We identified five reactions (A-E) among the four subnetworks related to chronic ethanol exposure (**Table 2**); all were FA hydrolysis or transfer reactions catalyzed by phospholipases and transferases. These results indicate that reaction C ([LPC] + [LPC] → [PC]) displayed dysregulation only in WT mice, regardless of sex. Reactions A ([PE] + [LPC] → [PC] + [LPE]), B ([LPC] ↔ [PC]), and D ([PC] + [DG]→ [LPC] + [TAG]) were present in all comparisons, except in IM WT (ethanol exposure vs. control in male WT mice), while reaction E ([LPC] + [LPE]→ [PC]) was exclusive to IF TLR4-KO (ethanol exposure vs. control in female TLR4-KO mice). These results suggested distinct enzymatic dysregulation in IM WT compared to the remaining conditions; however, all enzymes participate in glycerophospholipid metabolism. Indeed, some can activate phosphatidylcholine, forming intraluminal vesicles ^39^. Several studies have shown that decreasing phosphatidylcholine, cholesterol, and ceramide levels in microvesicular bodies could inhibit exosome release through the activation of lipidomic enzymes ^40,41^.

**Table 2.**
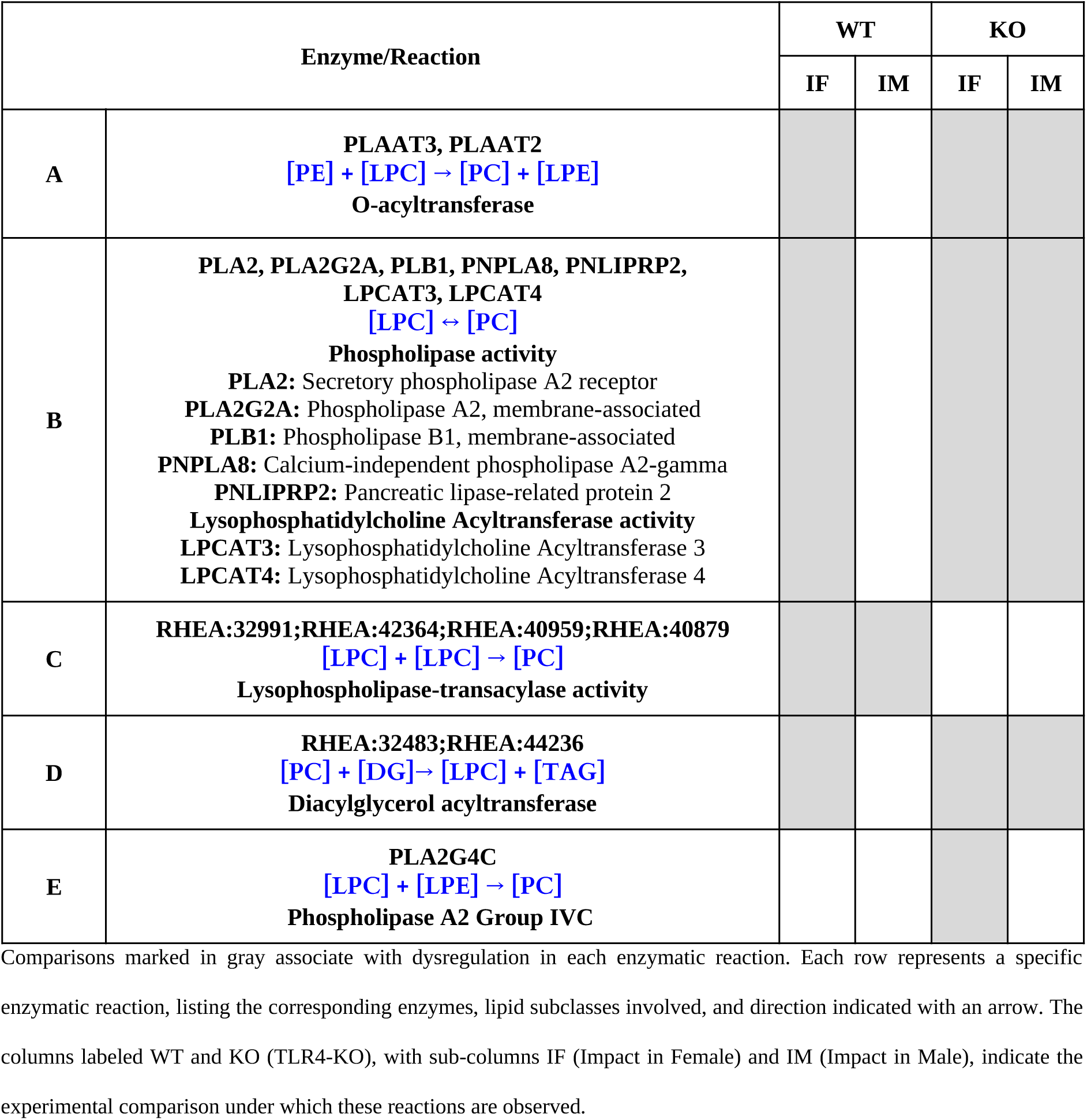
Enzyme reactions identified among the optimal subnetworks predicted by the LINEX^2^ enrichment algorithm in the four comparisons.

Activation of the TLR4 immune response and oxidative stress processes participate in chronic inflammatory diseases, including alcohol-induced neurological damage ^42^. Phospholipid oxidation, by modulating the activity of phospholipases such as PLA2, participates in ethanol-induced stress damage ^42,43^. For instance, PLA2 inhibitors prevent ethanol-induced neurodegeneration in rat organotypic brain slice cultures ^44,45^. Interestingly, our results demonstrated that PLA2 participates in reactions B and E (**Table 2**), suggesting PLA2’s involvement in the effects of ethanol exposure and regulation of the TLR4-associated immune response. Furthermore, reaction B belongs to Land’s Cycle (main process through which phospholipids are remodeled to modify their fatty acids composition), which interconnects with TLR4; these processes participate in lipid metabolism and inflammatory responses in cells, particularly in macrophages, and contribute to membrane remodeling processes related to permeability and secretory vesicle formation ^46,47^.

We also observed differences in Lipid Ontology (LION) enrichment analysis in our study when using lipids in the subnetwork as targets in the target list mode ( **Fig. 4C**). Focusing on terms related to lipid bilayer properties, such as "*transition temperature*", "*bilayer thickness*" and "*lateral diffusion*", we noticed notable variations between comparisons. We identified a low number of terms related to transition temperature and bilayer thickness in IM WT and a high number of terms related to transition temperature and bilayer thickness in IM TLR4-KO. We identified the terms “very low transition temperature” and “very low bilayer thickness” in IM WT and the terms “high transition temperature” and “high bilayer thickness” in IM TLR4-KO. Potential alterations in the lipid bilayer properties of plasma EVs suggest the existence of adaptive mechanisms in response to chronic ethanol exposure ^48^. Biochemical abnormalities in the brain of chronic heavy drinkers are associated with increased cell membrane rigidity and changes in phospholipid composition to compensate for brain structural changes detected by CT (computed tomography) and MRI (magnetic resonance imaging) in alcohol abusers ^49^. Lipid bilayer permeability influences how and when exosomes release their cargo into recipient cells, and while a reduction in membrane rigidity remains crucial for facilitating the formation of the highly curved vesicles essential for intracellular cargo trafficking ^50^, a higher bending rigidity generally contributes to increased vesicle stability, making it more difficult to deform or rupture ^39^.

### Differential Abundance of Lipid Species in EVs During Chronic Ethanol Exposure: Impact of TLR4, Sex, and Genotype

Given that each lipid class or subclass has specific functions, we conducted a differential abundance analysis to determine the effect of TLR4 on EV lipid distribution and composition during chronic ethanol exposure, focusing on differences by sex. We compared chronically ethanol-exposed mice with control animals in both sexes and genotypes; **Figure 5A** depicts the distribution of significant lipid species among the different lipid subclasses. We observed 16 significant lipid species in female WT (IF WT) and 45 in male WT mice (IM WT). Similarly, we observed 22 significant lipid species in female TLR4-KO mice (IF KO) and 119 in male TLR4-KO mice (IF KO). **Table S4** summarizes those lipids with significantly differential abundance following chronic ethanol exposure, categorized by LFC and comparison.

**Figure 5.**
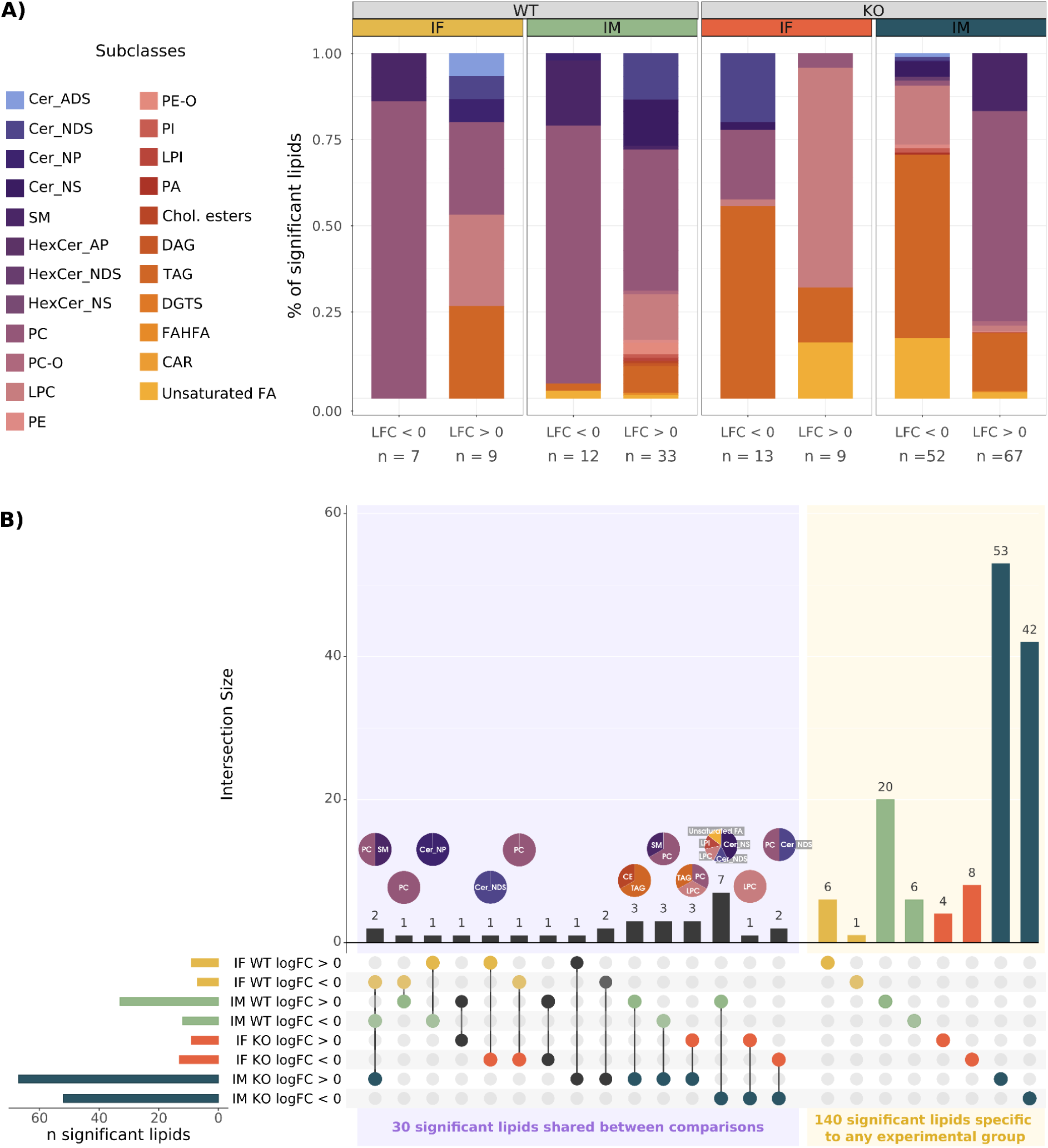
Summary of differential abundance analysis and molecular lipid profiles in extracellular vesicles for the four study comparisons. **(A)** Bar plots displaying significantly altered lipids with a p-value adjusted ≤ 0.05, separated by the log-fold change (LFC) and classified by subclass. **(B)** Upset plot illustrating the differential abundance analysis of lipids. The LFC sign distinguishes each comparison’s data. Horizontal bars represent the number of significant lipids in each comparison, color-coded for clarity. Vertical bars indicate lipids in the intersection of the groups, denoted with a colored dot underneath. A dot’s color indicates the comparison. Pie charts above the vertical bars show the lipid subclass of the intersecting lipids. Comparisons: ethanol exposure vs. control in female WT mice (IF WT), ethanol exposure vs. control in male WT mice (IM WT), ethanol exposure vs. control in female TLR4-KO mice (IF KO), and ethanol exposure vs. control in male TLR4-KO mice (IM KO).

Analysis of the lipid subclasses to which significant lipids belong revealed significant differences between comparisons (**Fig. 5A**). Chronic ethanol exposure caused remodeling of PC species in WT animals, decreasing the PC/LPC ratio in both males and females. Although a significant number of PC species increased in chronically ethanol-exposed mice, most identified PC species displayed decreased abundance; meanwhile, LPC species increased in chronically ethanol-exposed mice of both sexes. In TLR4-KO animals, we observed a significant differential abundance of these glycerophospholipids between males and females. Significantly, TLR4-KO males displayed an opposite pattern of differential abundance of these lipid species compared to their WT counterparts. A more significant number of LPC species had a lower differential abundance, and the number of PC species with a higher differential abundance was particularly significant, increasing the PC/LPC ratio; thus, the proportion of PC/LPC differed across the conditions. Notably, LPC (synthesis and catabolism mainly taking place in the liver) is considered a potential biomarker for liver diseases since numerous liver pathologies have encountered alterations in plasma LPC levels ^51,52^. Specifically, the PC/LPC ratio has been implicated in immune responses, particularly in disease-associated inflammation ^53^; for instance, LPC modulated immune signaling, including a role in TLR2/1- and TLR4-mediated signaling pathways ^54^.

Chronically ethanol-exposed WT and control TLR4-KO male mice displayed higher amounts of significant lipid subclasses than the remaining groups (**Fig. 5A**); moreover, both these groups shared specific lipids (e.g., PI and some Ceramides). Further analysis of differences in male mice revealed the higher abundance in lipid subclasses belonging to the Sphingolipids superclass in control WT mice and chronically ethanol-exposed TLR4-KO mice. PI has been associated with inflammatory responses in immune cells ^55^, while ceramides and sphingolipids participate in EV biogenesis and secretion ^1,56,57^; moreover, TLR4 can directly impact Ceramides, SM (main class Sphingolipids), and TAG ^58–60^. We also observed differences according to mouse genotype independent of sex (**Fig. 5A**). Chronically ethanol-exposed male and female WT mice displayed a significant increase in Ceramides while SM decreased, whereas the abundance of these lipid species demonstrated a reversed pattern in TLR4-KO mice. While Ceramides displayed a lesser degree of differential abundance, SM exhibited a greater differential abundance in ethanol-exposed TLR4-KO mice (**Fig. 5A**), with the increase in Ceramides coming at the expense of SM ^61^. For instance, Cer_NDS d36:1 increases and SM d36:1 decreases in female WT mice (**Table S4**), and Cer_NDS d36:1 also decreases in female TLR4-KO mice. We observed similar patterns in males; certain Ceramide species increased in WT mice but decreased in TLR4-KO animals, and some SM (e.g., SM d41:1 or SM d42:1) decreased and increased in WT and TLR4-KO mice, respectively. Thus, our study confirms the role of the TLR4 in Ceramide accumulation during chronic alcohol exposure and supports the dysregulation observed in other human and animal models studies ^62^. TAG levels decreased in both male and female TLR4-KO mice (**Fig. 5A**). TLR4-KO mice do not accumulate TAG during alcohol consumption, thereby preventing alcoholic fatty liver disease ^63^.

We also identified sex-specific EV lipid species after analyzing the number of significantly altered lipids shared between the different comparisons after ethanol exposure (**Fig. 5B**). Specifically, we identified 7 and 26 specific lipid species in female and male WT mice, as well as 20 and 95 specific lipid species in TLR4-KO female and male mice, respectively. Moreover, certain species shared between comparisons displayed opposite patterns of abundance. After chronic alcohol exposure, the cerebral cortex of ALDH2-deficient mice had more significant changes in the composition and content of lipid classes than WT mice, affecting structures (e.g., the blood–brain barrier) and aggravating neuroinflammation ^64^. In our case, the higher content of significant lipid species in TLR4-KO male mice could relate to the restoration of inflammatory processes since we previously reported that chronically ethanol-exposed TLR4-KO mice possessed protection from neuroinflammation and brain damage ^12,65,66^.

These results also suggest that the variables - ethanol exposure, sex, and genotype - associate with changes in the lipidome, demonstrating their involvement in lipid-associated tissue homeostasis or lipid metabolism disorders ^1^. Perpiña-Clerigues et al. also demonstrated sex and genotype differences induced by binge ethanol exposure in adolescent mice ^21^. Here, we identified the same changes in the EV lipidome previously associated with specific pathophysiological processes in chronic alcohol consumption; importantly, we found that TLR4 directly affects lipidomic changes, confirming TLR4 as a potential treatment target.

### Limitations and considerations

As relatively new fields, EV analysis and lipidomics remain at the forefront of innovation but have certain limitations. This study aimed to provide data on individual lipid species to enhance lipidomic pathway analysis, acknowledging that lipid species within the same class can have distinct biological functions; however, the lack of standardization in lipid nomenclature and integration into computational tools represents a significant limitation ^67^. While lipid bioinformatics packages such as LINEX^2^ offer new perspectives on metabolism, they primarily highlight differences between subtract-product reactions in distinct groups and do not address the direction of enzyme-catalyzed reactions. Additionally, although the EVs used in this study resemble exosomes in size and shape, we have referred to all species as EVs due to uncertainty regarding their identification.

## Conclusion

Our results demonstrate the involvement of TLR4 and sex in inducing alterations in the EV lipid metabolic networks generated by LINEX^2^ and LION enrichment analysis and the abundance of EV lipids induced by chronic ethanol exposure in female and male mice. Interrogating lipid abundance with this sensitive lipidomic-based strategy provides deeper insight into the molecular mechanisms underlying biological processes associated with sex, alcohol consumption, and the TLR4-mediated immune response. This approach opens a new avenue for biomarker identification and therapeutic development or as an intervention route.

## Declarations

### Ethics approval and consent to participate

All the animal procedures were carried out in accordance with the guidelines approved by the European Communities Council Directive (2010/63/ECC) and Spanish Royal Decree 53/2013, modified by Spanish Royal Decree 1386/2018, and were approved by the Ethical Committee of Animal Experimentation of the CIPF (Valencia, Spain).

### Availability of data and materials

The datasets generated and analyzed during the current study and programming scripts are available in the GitHub repository: https://github.com/carlapercle/lipidomics_chronics_mice.git.

### Authors’ contributions

CPC analyzed the data; MP and FGG designed and supervised the bioinformatics analysis; SM obtained mouse plasma samples and isolated EVs from plasma; CPC, MP, FGG, and CGR helped in the interpretation of the results; CPC, MP, FGG, and CGR wrote the manuscript; CPC, CGR, MP, and FGG writing-review and editing; MP and FGG conceived the work. All authors read and approved the final manuscript.

## Acknowledgments

The authors thank the Principe Felipe Research Center (CIPF) for providing access to the cluster, co-funded by European Regional Development Funds (FEDER) in the Valencian Community 2014-2020. The authors also thank the Genomics and Proteomics Unit at the University of Alicante, the Electron Microscopy Service at the Príncipe Felipe Research Centre, and Stuart P. Atkinson for reviewing the manuscript.

